# A Computational Account of Real-World Attentional Allocation Based on Visual Gain Fields

**DOI:** 10.1101/2023.09.18.558354

**Authors:** William J Harrison, Imogen Stead, Thomas SA Wallis, Peter J Bex, Jason B Mattingley

## Abstract

Coordination of goal-directed behaviour depends on the brain’s ability to recover the locations of relevant objects in the world. In humans, the visual system encodes the spatial organisation of sensory inputs, but neurons in early visual areas map objects according to their retinal positions, rather than where they are in the world. How the brain computes world-referenced spatial information across eye movements has been widely researched and debated. Here we tested whether shifts of covert attention are sufficiently precise in space and time to track an object’s real-world location across eye movements. We found that observers’ attentional selectivity is remarkably precise, and is barely perturbed by the execution of saccades. Inspired by recent neurophysiological discoveries, we developed an observer model that rapidly estimates the real-world locations of objects and allocates attention within this reference frame. The model recapitulates the human data and provides a parsimonious explanation for previously reported phenomena in which observers allocate attention to task-irrelevant locations across eye movements. Our findings reveal that visual attention operates in real-world coordinates, which can be computed rapidly at the earliest stages of cortical processing.

## Introduction

The behaviour of any animal is shaped by the sensory information it extracts from the environment. All sensory systems, however, only encode a finite amount of information. In the human visual system, for example, only a small portion of central vision is capable of processing information in high resolution. We therefore need to actively construct a high-resolution representation of a scene over time by coordinating saccadic eye movements with shifts of visual attention ^1^. A shift of attention to a location in peripheral vision prioritises processing of objects appearing at that location, while a saccade shifts the high-acuity fovea directly to an object. Such shifts can occur multiple times per second in naturalistic viewing conditions ^2^, and can improve discriminability of attended or fixated objects ^3^. The coordination of eye movements and spatial attention is thus fundamental to our ability to represent richly detailed information despite the information processing limits of sensory systems.

Despite decades of psychophysical and neurophysiological experimentation, there is no consensus on how the brain coordinates saccades and shifts of attention. While a saccade target is brought to the fovea after an eye movement, objects throughout the rest of the visual field are abruptly shifted to new retinal locations. Saccades also disrupt visual processing by reducing contrast sensitivity ^4–7^. Interestingly, however, these large-scale changes do not seem to interrupt our subjective visual experience, and we maintain phenomenal *visual stability* across saccades. One hypothesis is that visual stability is achieved by trans-saccadic updating of attentional “pointers” that track the locations of objects of interest ^8^. On this hypothesis, neural activity throughout oculomotor brain areas serves to predict the consequences of eye movements, but not by predicting the impending retinal image per se. Instead, the prediction is a shift in the focus of attention such that, at the completion of the saccade, prioritised processing is already directed to the new locations of attended objects ^9^. Various forms of this attentional remapping hypothesis have been proposed ^10–12^. While broadly similar, these hypotheses differ in how they seek to reconcile the psychophysical data with results from relevant neurophysiological investigations.

A recently advanced account is that visual stability begins in the first stages of visual processing ^13^. It is well known that neurons in primary visual cortex (V1) encode basic features of visual objects in an eye-centred frame of reference. These neurons are selective for small regions of oriented contrast ^14^ that form the building blocks for all visual function ^15^. A large proportion of V1 neurons also encodes gaze position ^16,17^. These so-called “gain-field neurons” not only encode an object’s location in an eye-centred frame of reference, they also represent where the eyes are directed in external space. A recent study demonstrated that these pieces of information are sufficient to recover the world-referenced location of visual objects. Moreover, this information can be decoded from V1 neurons in close to real time, even across eye movements ^13^. Rather than being supported by attentional pointers, therefore, visual stability may be computed early in the stream of processing, allowing attention to be allocated to objects’ true locations. This proposal is appealing because the speed and accuracy of such gain-field calculations allows for an almost continuous percept across saccades, but this model has not been tested with psychophysical data.

Here we investigated trans-saccadic changes in attentional selectivity with unprecedented spatio-temporal precision. Our primary aim was to test the accuracy of attentional selectivity in tracking an object’s position in the real world during eye movements, and to construct a model that calculates the real-world locations of objects and directs attention accordingly. We therefore combined psychophysical measurements of orientation-tuned contrast sensitivity, eye tracking, and dynamic classification image analysis. We used a novel classification image paradigm in which visual noise was added across the display, allowing us to probe attentional selectivity at arbitrarily specified locations using reverse correlation ^18–20^. By quantifying observers’ attentional template across space and time, our approach characterises the surprising precision with which observers maintain attentional selectivity across saccadic eye movements. We further developed an ideal observer model that allocates attention in world-referenced coordinates as computed by idealised gain-field neurons, and we show that this model explains not only our main findings but also previously reported influences of attention at non-cued locations ^e.g. 21^.

## Results

### Overview

Seven observers each completed 12,000 trials of a dual task in which they were required to identify a target at one location, while making a goal-directed saccadic eye movement to another location. As shown in Figure 1 (a,b), the display was designed so that the eye movement task mimicked a relatively natural sequence of horizontal and vertical eye movements across trials. Observers were required to maintain fixation in the centre of a green box, which, after a random delay within each trial, jumped to a new screen position (Figure 1c). Observers were instructed to make a saccadic eye movement as quickly and accurately as possible to the new location of the green box. Importantly, prior to the start of each trial, a visual cue (red box in Figure 1) indicated the upcoming perceptual target location with 100% validity. The target duration was brief (17ms), and observers were instructed that it could appear at any time during a trial. The certainty with which the cue indicated the target location, in addition to the uncertainty in target onset time, motivated observers to attend to the target location for the duration of the trial, and therefore across the saccade. At the end of each trial, observers reported whether a light or dark bar had appeared at the cued location. Target amplitude was controlled by an adaptive staircase, and was embedded in visual noise, as described in detail below.

**Figure 1.**
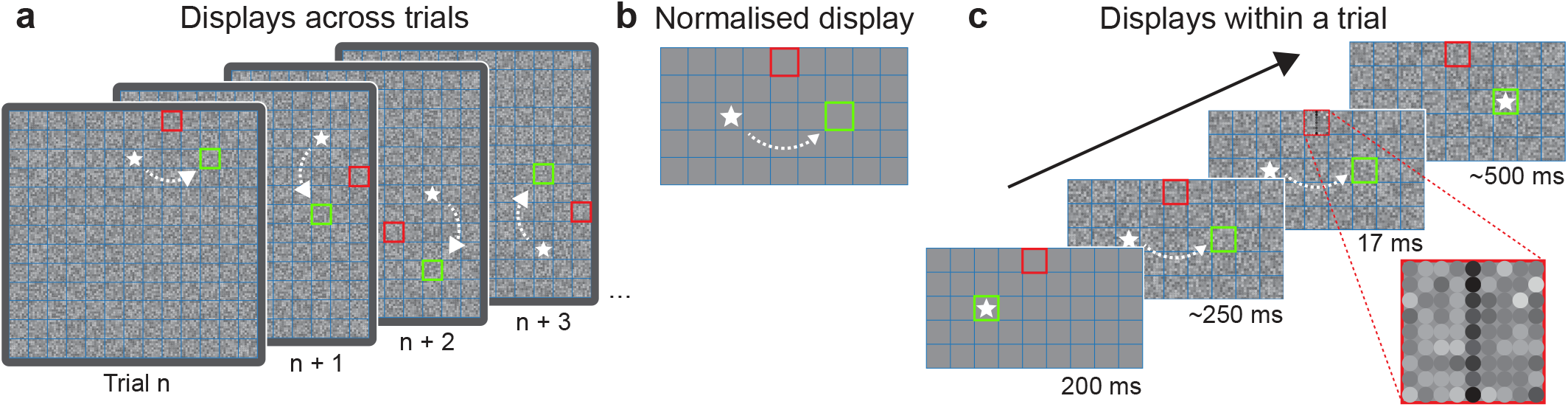
Experimental design and displays used to measure trans-saccadic attentional updating. Observers performed a dual task in which they were required to make a goal-directed saccade to one location while monitoring for a perceptual target at another location. a) Example displays across trials. The full display extended approximately 30° x 30°, and was divided into a 13 x 13 grid. Within each example trial display in this figure, the white star (not present in the actual displays) represents a hypothetical observer’s fixation position at the start of the trial, the green box indicates the saccade goal, the arrow indicates the direction of the required saccade, and the red box indicates the perceptual target location. Note that the initial fixation position of each new trial was the same as the saccade goal on the previous trial, which elicited a relatively natural progression of eye movements across trials. The required saccade direction (up, down, left or right) was selected pseudo-randomly from trial-to-trial and had an amplitude of 4 grid locations (9°). b) Prior to analyses, the differential organisation of displays across trials was normalised to a common reference frame. It can be appreciated in this normalised display that the perceptual target was presented at the same eccentricity (6.4°) from the initial fixation location and the required saccade target. c) Progression of displays within a trial. Observers began each trial by fixating the centre of the green box. Prior to the start of the trial, the perceptual cue was shown for 200ms, cueing the observer to covertly attend to this location. After an additional random interval, the green fixation box shifted to a new location, indicating the saccade goal. The perceptual target, shown here as a relatively dark bar, had a duration of 17ms and could appear at any time during a trial with equal probability. Perceptual reports (“light” or “dark”) were made at the end of each trial. Throughout each trial, dynamic visual noise was displayed at a rate of 60Hz. Noise was organised as 9 x 9 spots (0.25° diameter) within each grid location. The luminance of each spot of noise was randomly drawn from a Gaussian distribution, with a mean contrast of 12.5%.

### Data pre-processing

To investigate changes in perception across eye movements, each trial was binned according to when the perceptual target was displayed relative to the onset of the saccade (bin width = 17ms). Prior to analyses, data were screened to include only trials in which the eye movement task was completed successfully (see Methods). In brief, we included trials in which saccadic latencies were between 100ms and 500ms relative to the onset of the saccade goal (Figure 2a). We excluded trials in which an observer’s fixation deviated by more than 2° from the centre of the required initial fixation box prior to saccade onset, and trials in which the saccade endpoint deviated by more than 25% of the required saccade amplitude from the saccade goal (Figure 2b). Importantly, after all pre-processing steps, we retained hundreds of trials per saccade-locked time bin per participant, as shown in Figure 2c.

**Figure 2.**
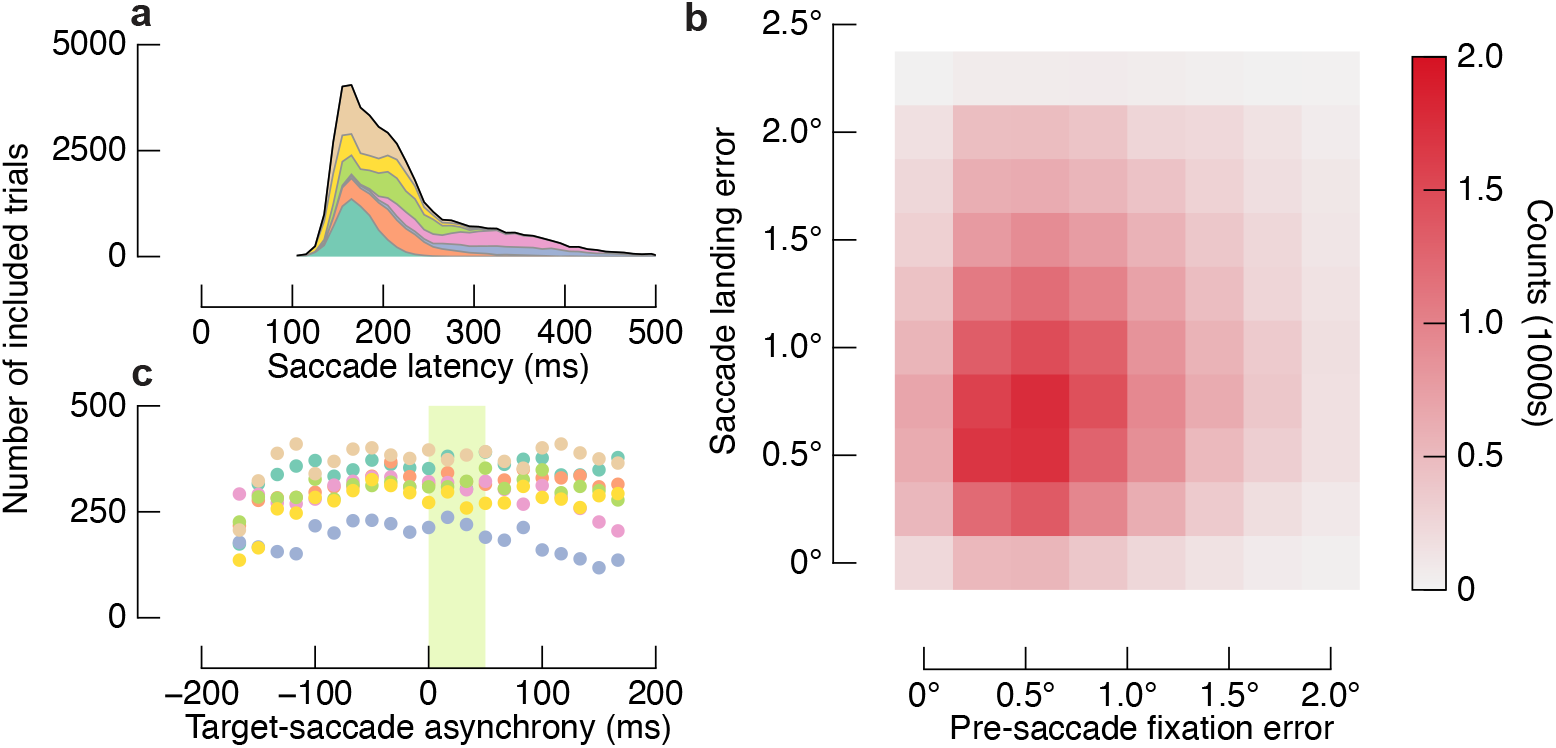
Eye movement related parameters. a) Stacked distributions of saccadic latencies uniquely coloured for each observer (N=7). We excluded trials in which saccadic latencies were shorter or longer than 100ms or 500ms, respectively. b) Fixation errors for all included trials. Pre-saccade fixation error and saccade landing error are shown as joint-distributions, with bin-counts indicated as per the colour bar. Trials were excluded if pre-saccadic fixation deviated by more than 2° from the initial fixation box, or if the saccade landed more than 2.3° from the centre of the saccade target (i.e. if saccade landing error was more than 25% of saccade amplitude). Other exclusion criteria are described in the Methods. c) Number of included trials in each time bin. Colours show different participants, as per (a). We retained hundreds of trials per time bin per participant.

### Trans-saccadic perceptual performance

The target contrast was adaptively controlled from trial to trial to maintain each observer’s overall accuracy at approximately 76%. The perceptual task was therefore relatively difficult, requiring attention to be successfully allocated to the cued target location. The brief duration and uniformly random onset timing of the target allowed us to measure perceptual performance before, during and after a saccade with higher temporal precision than is typically measured ^9,22^.

Figure 3a shows mean accuracy in the perceptual task as a function of target onset time relative to saccade onset time. Coloured data in the figure show each individual’s accuracy, and the black line represents the mean for the group. Negative values on the x-axis indicate targets presented prior to the eye movement, and the green shaded region spanning from 0ms to 50ms indicates the modal duration of saccades across observers. Although accuracy dropped to chance (i.e. proportion correct = 0.5) during the saccade itself, observers’ accuracy returned to above-chance performance by the completion of the saccade (i.e. by time bin +50ms). Mean performance was slightly lower after the saccade than before it, but this difference was driven by two participants. Generally, accuracy across observers was very similar due to the adaptive staircase.

**Figure 3.**
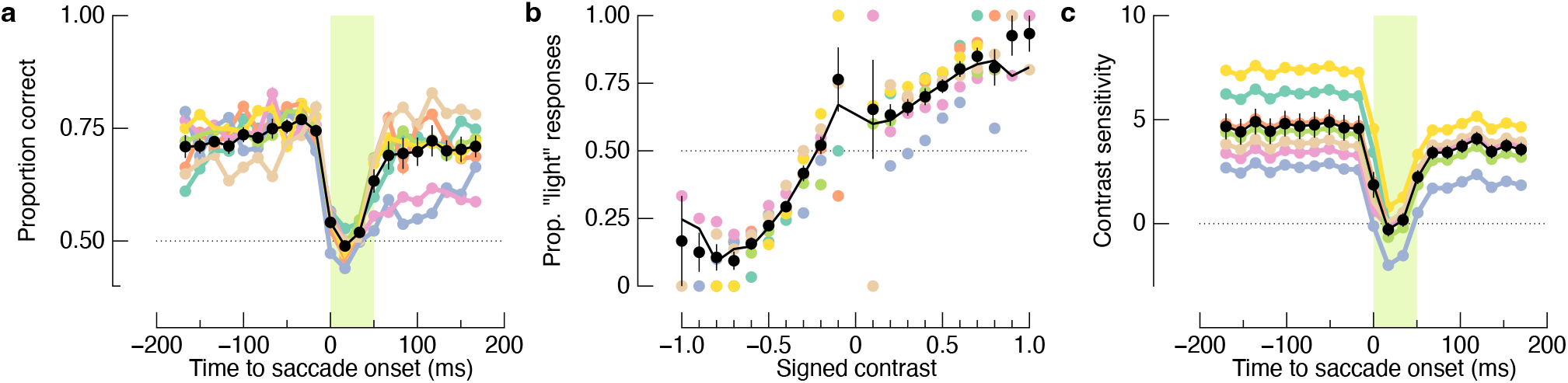
Perceptual accuracy and contrast sensitivity across eye movements. a) Proportion of correct target identifications as a function of target onset relative to saccade onset. Group means are shown as black symbols, and colours indicate different observers. Despite the difficulty of the perceptual task, performance dropped only during the saccade itself, which is indicated by the green shaded region. b) To assess the impact of saccades on contrast sensitivity, we measured perceptual reports for a range of target contrasts. Data within each time bin were fit with a GLM (black line), the slope of which quantifies contrast sensitivity. The model fit shown here is for data averaged across observers, but in the main analyses we fit the model to each observer’s data separately. c) Contrast sensitivity is affected by eye movements only during the saccade itself. Error bars in all panels show ± one standard error across observers, which is smaller than the point size in many cases.

As shown in Figure 3b, we measured perceptual reports across a range of target contrasts. We modelled contrast sensitivity at the attended target location with a generalised linear model (GLM), where the slope of the model indicates sensitivity to contrast (see Methods). We fit a multilevel GLM (GLMM) to the data, modelling different time bins as a random effect to derive trans-saccadic contrast sensitivity functions for observers, who were also included in the model as a random effect. Figure 3c shows that contrast sensitivity fell to zero during the intra-saccadic period, but was restored within 17ms of the end of the modal saccade offset.

The behavioural data reported above show that perceptual performance was impacted by eye movements only when the target presentation overlapped with the saccade itself. The intra-saccadic loss of sensitivity is likely due to the target’s retinal motion, which necessarily diminishes the target’s effective contrast ^4^. Because the attentional demands of the perceptual task were maintained with an adaptive staircase, the rapid recovery of perceptual performance immediately following the saccade indicates that observers were attending to the target location within 17ms of the completion of the saccade. Such temporally precise attentional control cannot be explained by stereotyped behaviour that developed across many repeated trials because the direction of the impending saccade was not known to the observer at the beginning of the trial.

### Dynamic orientation selectivity across saccades

We quantified dynamic changes in orientation-tuning across saccades using a reverse-correlation approach ^18,19,23,24^. As with the behavioural data above, we first time-locked each frame of the dynamic visual noise to saccade onset on a trial-by-trial basis. We then performed a spectral analysis at each time point using biologically-inspired filters ^15^, quantifying the orientation and scale of various random patterns within the noise after removing the target (see Figure 4). Using a series of GLMMs, we weighted the response of each filter to predict observers’ perceptual reports. The resulting three-dimensional sensitivity kernels describe observers’ tuning to orientation and spatial frequency at each time point.

**Figure 4.**
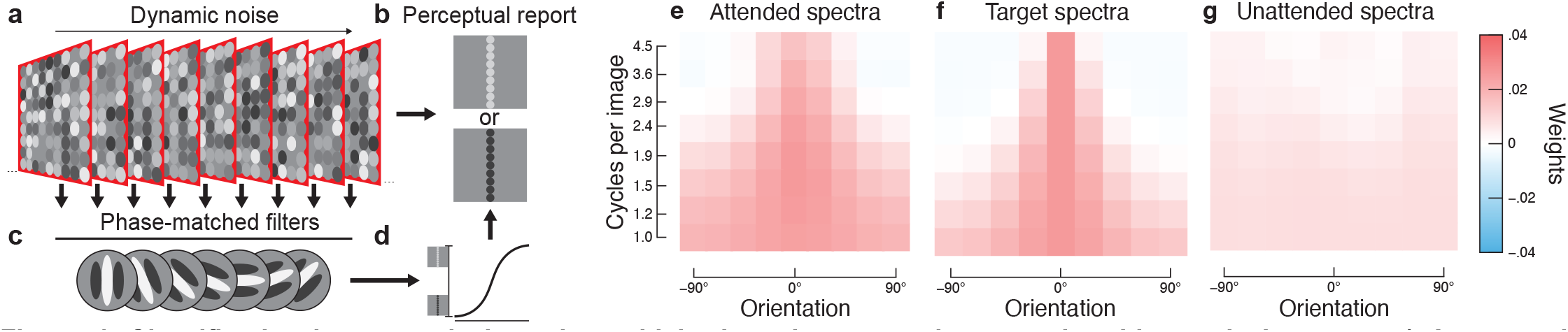
Classification image analysis and sensitivity kernels averaged across time bins and observers. a) An example sequence of dynamic noise presented at the attended perceptual target location. Each noise spot was randomly sampled from a Gaussian distribution (12.5% contrast) at a rate of 60Hz. Note that at some point during the sequence of frames, the retinal projection of the noise frame changed when an observer made a saccade from one screen location to another (see Figure 1c). b) At the end of each sequence of noise, observers reported whether they saw a relatively light or dark bar. The goal of the classification image analysis is to predict perceptual reports from the preceding dynamic visual noise. c) Spatial patterns are detected in each noise frame by applying a bank of oriented filters of various spatial scales, phase-matched to the perceptual target. The strength of the response of a given filter is determined by the extent to which random spatial patterns of noise match the tuning of the filter. Prior to this analysis, targets were removed from the noise. d) Perceptual reports were modelled by weighting each filter output at each time point before, during and after a saccade. e) Orientation and spatial frequency tuning at the cued perceptual target location. Data are marginalised over time. Warmer colours indicate stronger sensitivity to those features, as indicated by the colour bar at the right. f) Orientation and spatial frequency spectra for a noise-free target. We applied the same filters as used in the reverse correlation analysis to a single target, and then standardised the filter outputs for comparison with the sensitivity kernel shown in (e). g) No tuning was observed at a control location. The control location was a non-cued grid location, matched with the cued perceptual target location in all other aspects.

We first applied our reverse correlation analysis to visual noise presented at the attended perceptual target location (i.e. the red square in Figure 1). Figure 4e shows orientation and spatial frequency sensitivity kernels, collapsed across time bins and observers. There is clear orientation selectivity, with greater sensitivity to the target orientation (normalised to 0°) than to orthogonal orientations. The sensitivity to 0°-tuned filters is broadband, with relatively even weights regardless of frequency (cycles per image). We can see how closely observers’ tuning matched the properties of the perceptual target by analysing the spectral content of a noise-free target (Figure 4f). The observers’ tuning and the target spectra are in close agreement. The most noticeable differences are in the higher frequency channels, likely because of observers’ acuity limitations, but even these differences are small. Changes to filter parameters such as bandwidth, number, or spacing did not substantially alter these results.

We next performed the reverse correlation analysis on visual noise presented at a control location that was relevant for neither the perceptual task nor the eye movement task. The control location was matched with the perceptual target in all ways but was not cued as an upcoming target location. As shown in Figure 4g, there is no evidence of tuning at the control location, demonstrating that the orientation tuning in Figure 4e results from observers’ attentional allocation to the perceptual target location.

We next recovered the temporal dynamics of orientation tuning across saccades by performing the classification image analyses at each time point relative to saccade onset. The resulting spatio-temporal sensitivity kernels are shown in Figure 5. Figure 5a shows tuning at each time point relative to saccade onset for the spatial frequency band of 1.9 cycles per image. The band of warm colours centred on 0° and extending from the pre-saccade to post-saccade period is clear evidence of orientation tuning throughout the execution of the eye movement. Indeed, the marginal distribution shown in Figure 5b reveals an orientation tuning curve centred on the target orientation. Figure 5c shows three time points around the onset and offset of the saccade. There is clear tuning immediately before the saccade (“pre-sac”), at the offset of the saccade (“sac offset”) and 34ms following saccade offset (“post-sac”). Similar results were found for all spatial scales (see Supp Fig S 1). There were no statistically significant sensitivity kernels immediately following the saccade (while the eyes were in flight) at any spatial scale.

**Figure 5.**
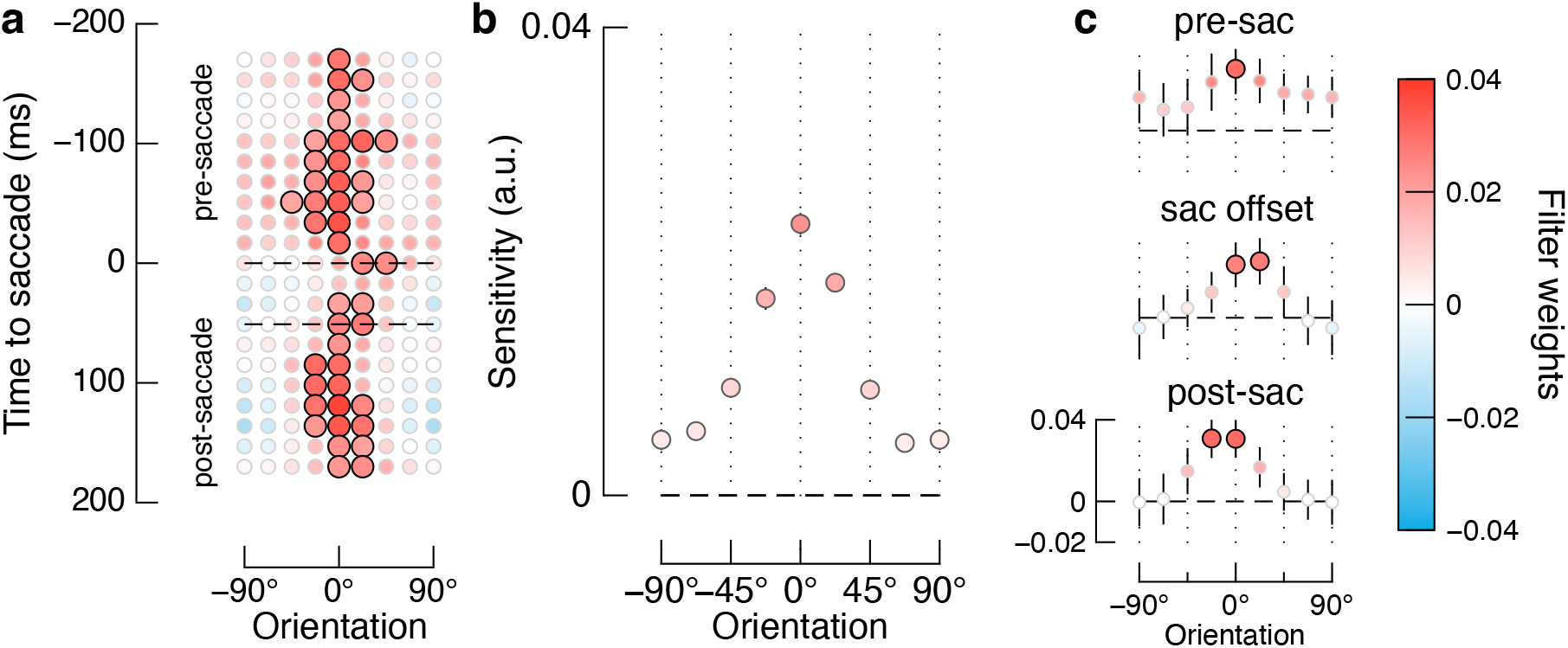
Orientation tuning at the attended location is sustained across saccades. a) Spatio-temporal sensitivity kernels for filters with a peak spatial frequency of 1.9 cycles per image. Data are time-locked to saccade onset, such that the dashed lines at 0ms and 50ms indicate sensitivity at saccade onset and modal saccade offset, respectively. Black outlines show statistically significant kernels (p < 0.05, uncorrected). b) Orientation sensitivity marginalised over time. Observers’ sensitivity is clearly tuned to the target orientation (0°). c) Sensitivity at saccade onset (“pre-sac”, top), at modal saccadic offset (“sac offset”, middle), and immediately after the saccade (“post-sac”, bottom). These slices show only a modest reshaping of the tuning curve across the saccade. Data are shown for the responses of filters with a peak spatial frequency of 1.9 cycles per image, but other frequency channels showed similar tuning (Supplemental Figure S1).

### Attentional allocation to non-cued locations

The analyses presented thus far reveal that observers’ attentional allocation is tightly locked on the cued target location with only a modest loss of tuning during the intra-saccadic period (Figure 5). These data suggest that attentional allocation is rapidly updated across the saccade following a time course that is well matched to the saccade dynamics. We can test a number of hypotheses regarding attentional allocation across eye movements by performing the same reverse correlation analyses as above but on noise presented at different grid locations relative to the saccade goal and perceptual target.

It has recently been proposed ^25^ that trans-saccadic attention involves two processes that are less temporally precise than the time course of the saccade: (1) a pre-saccadic shift of attention to anticipate the changed retinal position of the attended object, and (2) a post-saccadic withdrawal of attention from the no-longer relevant retinal location. Note that neither location is relevant for the saccadic or perceptual task in our experiments. Previous experiments that found evidence for attentional allocation at task-irrelevant locations did so by presenting perceptual probes at these locations, which might have biased observers’ attentional strategies. In contrast, in our study perceptual probes were only ever presented at the cued target location, and our analyses were performed on visual noise, thereby enabling us to recover attentional allocation without potentially biasing observers. We therefore tested for attentional allocation to either location – the pre-saccadic predicted cued location and the post-saccadic “attentional trace” location – before and after the saccade, by quantifying dynamic orientation selectivity at these locations. For the pre-saccadic predicted cued location, there was no tuning to the target orientation location at any time point. There was, however, broad tuning to the target orientation at the attentional trace location immediately post-saccade, but this tuning was not statistically significant. We discuss these predictions and data in more detail in the Supplemental Information.

### A generative observer model inspired by gain-field neurons

We developed a model observer motivated by a recent neurophysiological investigation of neurons in primary visual cortex (V1) of macaques ^13^. The visual receptive fields of V1 neurons are retinotopically organised and tuned to spatial frequency and orientation ^14^. A given V1 neuron therefore encodes the position of a visual feature in an eye-centred frame of reference. Importantly, however, populations of V1 neurons also encode the direction of gaze independently of visual stimulation. V1 neurons’ spike rates are systematically altered according to the current gaze position, a property referred to as the neurons’ ‘gain-fields’. The activity of V1 neurons is thus sufficient to uncover an object’s world-centred position, which can be computed by summing the position of the object in eye-centred coordinates with the current gaze position (see Figure 6a):

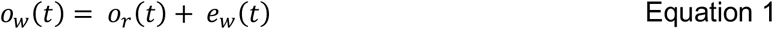

where *o*_*w*_(*t*) is the object’s real-world position at time *t, o*_*r*_(*t*) is the object’s position on the retina at time *t*, and *e*_*w*_(*t*) is the gaze position at time *t*. Morris and Krekelberg ^13^ demonstrated that the real-world position of a visual object can be decoded from the gain-fields of V1 neurons even across eye movements. The joint coding of retinal position and gaze position in V1 neurons therefore maintains a stable representation of visual objects in a real-world frame of reference. We propose that visual attention is allocated in this reference frame.

**Figure 6.**
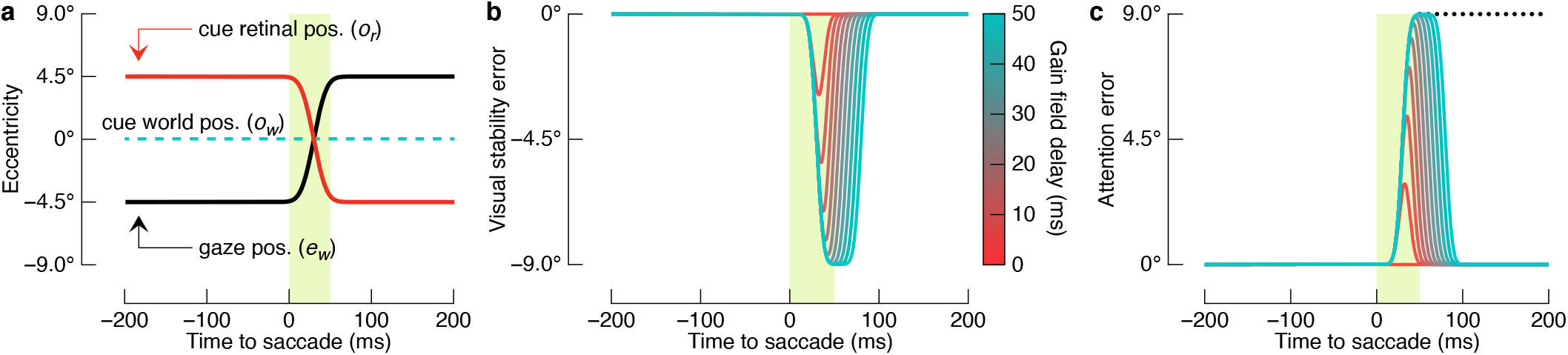
A model observer that allocates attention in real-world coordinates. a) Computing the cue’s true location as per gain-field neurons. In this example, the observer makes a 9° rightward saccade from -4.5° eccentricity to 4.5° eccentricity, while attending to the cue’s true location centred on the vertical midline of the display. Gain-field neurons encode the cue’s world location by jointly representing retinal position and gaze position, which, when summed, recover the true coordinates of the cue (Equation 1). The green shaded region is equivalent to the modal saccade duration in our experiment. b) Visual stability error as a function of the delay in gain-field neurons’ representation of gaze position. With zero delay, the cue’s true location is recovered perfectly across saccades. However, as the delay increases, so too does visual stability error up to a magnitude equivalent to the saccade amplitude. Note that at relatively high gain-field delays, the visual stability error persists even after the saccade is complete. Morris and Krekelberg ^13^ reported that V1 neurons in the macaque represent gaze position with a delay of 45ms. c) Error in attentional allocation as a function of gain-field delay. The transient mis-estimation of the cue’s true location after the saccade results in the erroneous allocation of attention. At the longest delays, attention is allocated to the retinotopic trace location (dotted line) immediately following the saccade.

We modelled gain-field computations in an uncertain template matching model ^26^ that performed the same dual task as human observers. The model executes an eye movement on each trial and then reports whether it detected a light or dark target bar in dynamic visual noise. As with the human observers, the model observer did not know in which noise frame the target would appear, and was therefore temporally uncertain. It discriminates a target by first computing a template response at each time point:

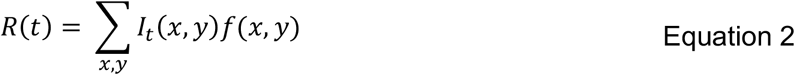

where *R*(*t*) is the template response at time *t, I*(*x, y*) is a 9 x 9 matrix of noise added to the display at time *t*, and *f*(*x, y*) is the template, which is identical in shape to a noiseless target. The model then reports whether a light or dark bar was presented, *δ*, by reporting the sign of the template response with the maximum absolute template response within a trial:

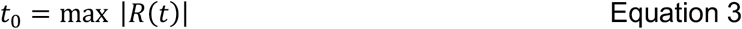

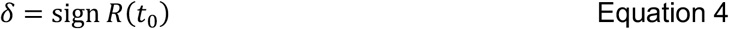

Given the temporal uncertainty inherent in the task, the model described with the equations above is an ideal observer model, i.e., it performs optimally given the noise in the stimulus. Critically, however, our model observer allocates attention in real-world coordinates. We modelled the allocation of attention by first estimating the real-world perceptual target location by summing the retinal position of the cued target location with the current gaze position as computed by idealised gain-field neurons ^13^ (Equation 1), and then applying the template to the noise at this location (Equation 2 - Equation 4).

We use the term *visual stability error* to refer to the difference between the cue’s real-world position and our V1 model’s estimate of the cue’s real-world position. If retinal and gaze position are encoded without noise and their summation is performed instantly, the cue’s real-world position is represented with no visual stability error (Figure 6a). The activity of gain-field neurons, however, is delayed with respect to the actual eye position, resulting in small but meaningful visual stability errors ^13^. Plotted in Figure 6b is a range of visual stability errors that can be expected from various delays in the encoding (or decoding) of eye position in our modelled gain-field neurons. As gain-field delay increases, so too does the visual stability error, which saturates around saccade offset before returning to zero. The model observer, which attends to the estimated real-world cue position, will thus transiently attend to non-cued locations when there is any delay in the gain-fields’ signalling of gaze position. As shown in Figure 6c, the error in attentional allocation can be as great as the saccade amplitude, shifting the focus of attention to the retinotopic trace location for a brief interval at the end of the saccade.

We analysed the model observer’s performance using the same reverse-correlation approach as for the human data. We estimated the gain-field delay by fitting the model output to the empirical data with a single free parameter representing the lag between the actual eye position and the estimated eye position. All eye movement dynamics were parameterised according to our empirical data, and other gain-field properties were as reported by Morris and Krekelberg ^13^ (see Methods). As shown in Figure 7a, the model observer’s spatio-temporal sensitivity kernels at the cued location were strongly tuned to orientation. Importantly, the model recapitulates the critical features of the empirical trans-saccadic spatio-temporal tuning functions (cf. Figure 5). The model’s spatio-temporal sensitivity kernels are generally more symmetric than the empirical data because the model is limited only by external noise. The estimated gain-field delay of our model observer was 15.7 ± 7.8ms, which is shorter than the 45ms delay found for real gain-field neurons in primary visual cortex ^13^.

**Figure 7.**
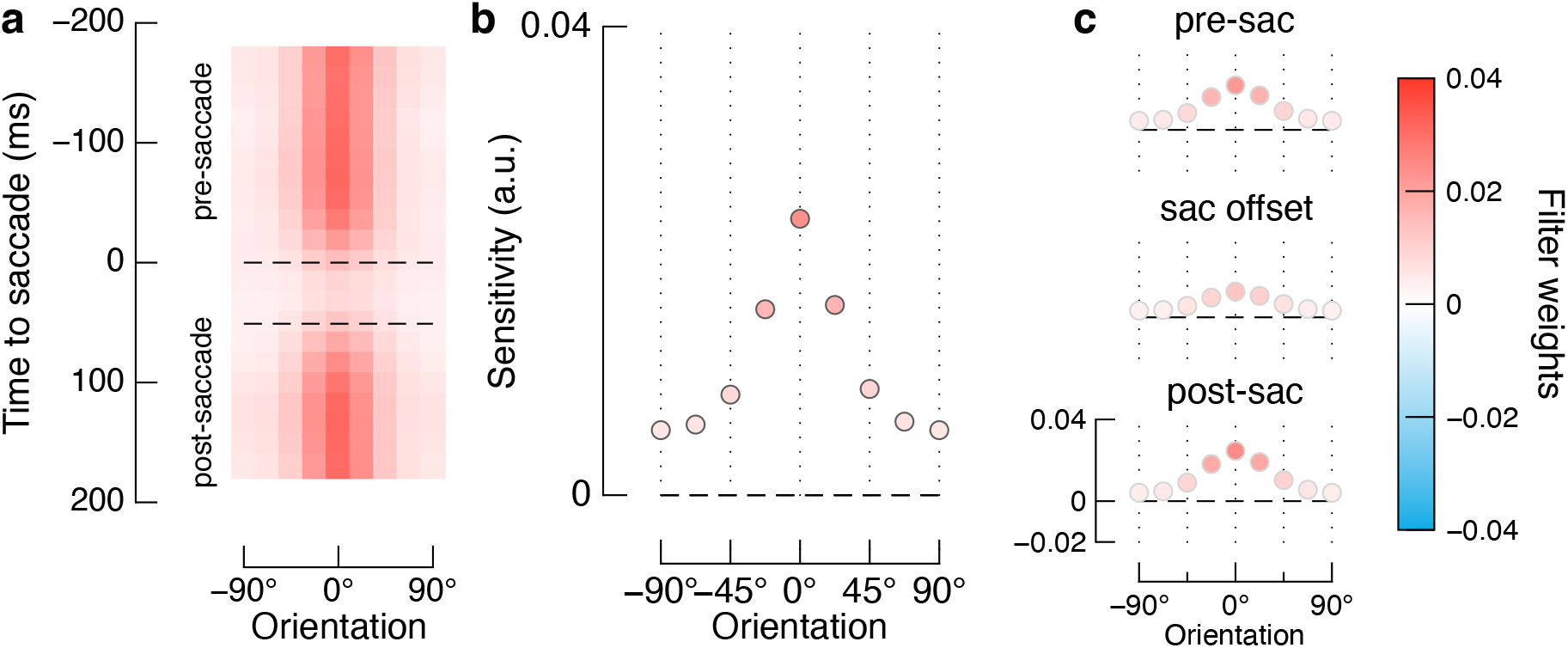
Modelled orientation sensitivity matches observers’ selectivity at both the cued and retinotopic trace locations. a-c) Spatio-temporal sensitivity kernels at the cued location, marginalised over spatial frequency. The model reproduces the empirical data shown in Figure 5.

## Discussion

We investigated observers’ ability to coordinate shifts of spatial attention with saccadic eye movements. We used a psychophysical approach in which observers made goal-directed saccades while monitoring for perceptual targets that were embedded in dynamic noise. By analysing the correspondence between the output of biologically constrained filters in response to the visual noise and observers’ perceptual reports, we were able to recover trans-saccadic orientation tuning functions for the first time. We found that both contrast sensitivity and orientation tuning are relatively unperturbed in the moments immediately preceding and following saccadic eye movements. Our data thus reveal that visual sensitivity can be maintained via coordinated shifts of attention and saccadic eye movements.

The coordination between attentional allocation and saccades is remarkably precise considering the typical time-course of voluntary and involuntary shifts of spatial attention in the absence of eye movements. After an eye movement has shifted an attentional cue to a new retinal location, we found a recovery of perceptual performance to pre-saccadic levels within 17ms for five out of seven observers, and within 33ms for all observers. However, following the presentation of a spatial cue like the one used in our experiment, performance benefits are not typically observed until at least 100ms after cue onset during steady fixation ^27,28^. The immediate recovery of post-saccadic contrast sensitivity we found cannot, therefore, be attributed to a de novo re-allocation of attention ^see also 22,29,30^. One increasingly popular explanation is that the focus of attention is remapped according to a prediction of the impending saccade ^8,9,31,32^. The predictive processes begin before saccade onset so that attention is allocated to the relevant location by the end of the saccade. However, a recent finding that the true location of a visual object is represented in the activity of neurons in primary visual cortex suggests that a predictive mechanism is not necessary to remap attention across saccades per se ^13^. Our model observer demonstrates that the computational burden of maintaining the focus of attention across eye movements can be simplified relative to hypotheses based on oculomotor prediction ^33^. Although there is no specific predictive mechanism in our model, the activity of gain-field neurons may nonetheless be modulated by predictive signals from somatosensory or oculomotor areas ^34^.

Our model observer is motivated by the recent finding that neurons in macaque primary visual cortex track the real-world positions of objects in close to real time, even across eye movements ^13^. The encoding of objects’ real-world locations at an early stage of visual processing provides an account of how objects can be tracked across eye movements with relatively little cognitive effort. Most people are unaware of where they have looked within just the last few seconds ^35,36^, even when they attend to the saccade target ^36^. This apparent disregard for the primary means by which we explore the visual world is impressive considering the extensive processes they are thought to invoke ^37^. The joint coding in early vision of an object’s basic features and the gaze position of the eyes greatly simplifies the coordination of cognitive operations that involve attentional allocation, working memory ^38^, and motor execution ^39^. By tracking real-world locations at the earliest stages of visual processing, higher cognitive processes can operate in a real-world frame of reference, rather than having to engage in additional computations to derive this information. Indeed, there is now evidence that the brain codes conceptual information in a low-dimensional format analogous to a world-centred representation of the environment ^40^.

Our gain-field model of attentional allocation provides a mechanistic account of attentional benefits at non-cued locations after saccades ^21,22,41^. Within a brief temporal window after saccade offset, attentional benefits have been found at the pre-saccadic retinal position of a target cue, even though this location no longer corresponds to the real-world position of the cue. Importantly, our model observer allocates attention in real-world coordinates calculated by gain-field neurons, rather than allocating attention in a retinotopic frame of reference that must be updated with each saccade ^c.f. 8^. When the delay between the current eye position and the gain-field calculation of eye position is sufficiently long, the model allocates attention transiently to the attentional trace location, as has been found for human observers in previous studies ^e.g. 21^. Our data thus reveal that a retinotopic attentional trace can be explained by a discrepancy between the current eye position and the estimated eye position, leading to attention being erroneously allocated when the eye movement ends (Figure 6). In principle, we could additionally model predictive pre-saccadic shifts of attention ^9^ as a corollary of the planned motor signal, as has been done previously ^33^. However, we found no such pre-saccadic effects in our data, consistent with the recent proposal that pre-saccadic shifts of attention depend on task demands ^42^.

Our study also provides clear evidence that attentional allocation mitigates the impact of saccades on contrast sensitivity ^4–7^. As expected, we found a loss of contrast sensitivity during the intra-saccadic period. Sensitivity was highly consistent until the onset of the saccade itself, however, and returned to pre-saccadic levels within 17ms after the saccade was complete. Other investigations using full-field displays have found that contrast sensitivity decreases well before an eye movement begins ^e.g. 4,6^. Although the pre-saccadic loss of sensitivity has been attributed to masking ^4^, we found no such effect in spite of the dynamic contrast of the visual noise in our experimental displays. We suggest instead that pre-saccadic sensitivity loss found in previous studies can be explained, at least in part, by the absence of positional certainty for the target location. It is well known that visual processing is prioritised at the saccade goal prior to saccade onset ^43,44^. Such a shift in processing may have made the identification of visual probes more difficult in previous experiments in which the upcoming perceptual target location was uncertain. In contrast to previous experiments, observers in our study were cued reliably to the precise target location, reducing positional uncertainty and maximising the benefit of attentional allocation. This critical role of attention in sustaining contrast sensitivity across eye movements also helps to explain why people do not experience a loss of sensitivity before and after saccades as previously suggested ^4,6^, because attended objects do not undergo suppression in the same way as unattended objects.

Although the visual system is constrained in its ability to process information, visual detail can nonetheless be acquired across a rapid succession of eye movements. Our study shows that the processing of visual detail is sustained across saccades at attended but non-fixated locations. A model observer that estimates the true locations of objects and then allocates attention in real-world coordinates allows efficient coordination of attention and saccadic eye movements.

## Supplemental Information

**Fig S1.**
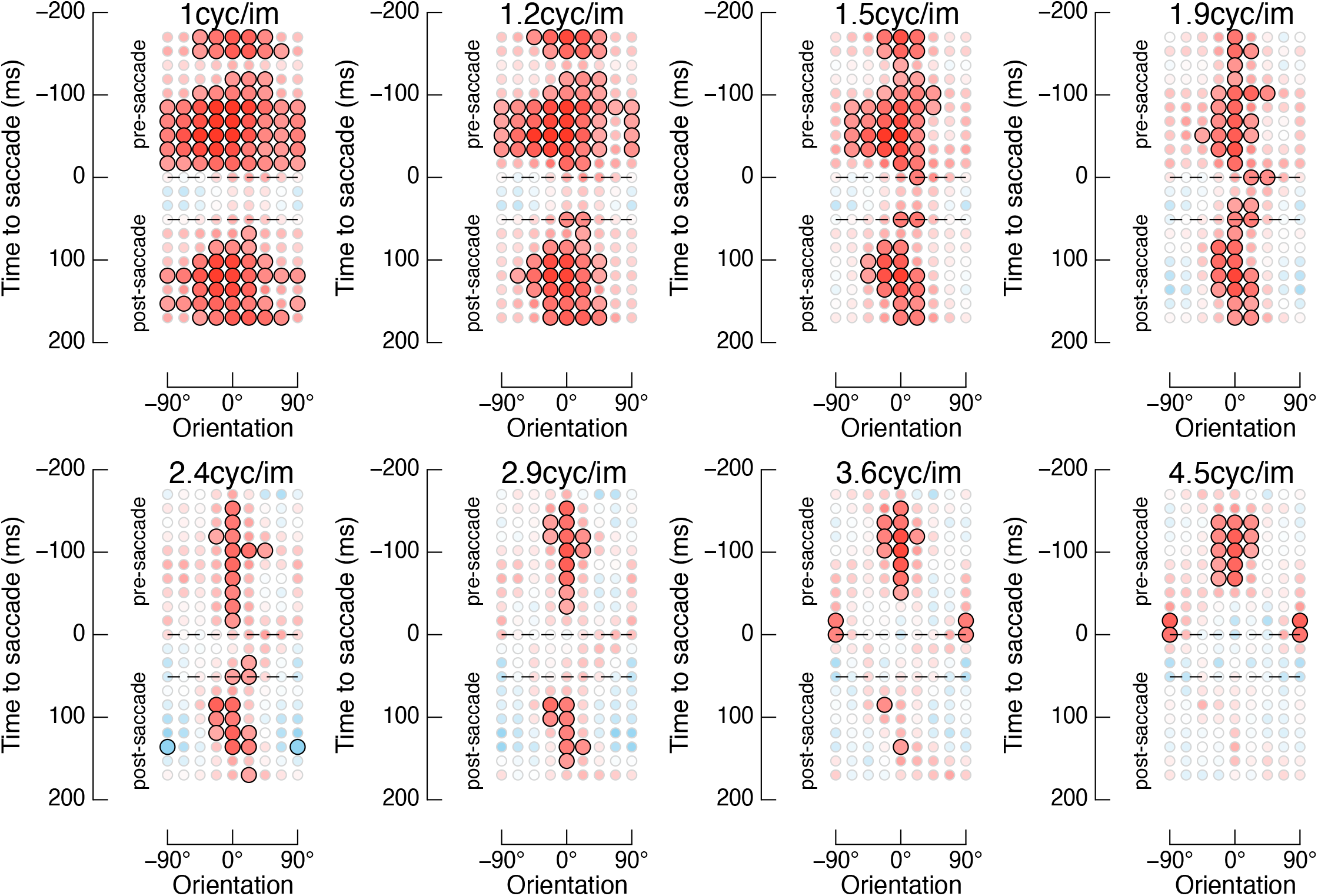
Orientation tuning at the attended location across different spatial frequency bands. The scale of the filter is shown at the top of each plot. Data are presented as in Figure 5. The range of significant kernels across orientations in the lower spatial frequencies is merely a result of the broad tuning of the filters: at these spatial scales, the responses of the filters are highly correlated across orientations.

### Post-saccadic attentional trace

Previous studies have found that, immediately *after* a saccade, attention remains allocated to the pre-saccadic retinotopic location of an attentional cue (Fig S 2a). This so-called *attentional trace* is often interpreted as evidence that visual attention is allocated in a retinotopic frame of reference ^21,22,41^. However, each study reporting evidence of an attentional trace has only been able to assess attentional allocation by presenting a perceptual target at the location of the “retinotopic trace”. This work, therefore, has been unable to rule out the possibility that observers’ attentional allocation across saccades is altered when they must respond to targets presented at the retinotopic trace location. Our study addresses this shortcoming: observers knew the target location with 100% certainty on each trial, but we were nonetheless able to probe attentional allocation at any screen location via reverse correlation of the visual noise. We tested for evidence of an attentional trace by quantifying orientation tuning at the retinotopic trace location. We omitted from the analysis data from the pre-saccade interval, because the retinotopic trace location only exists, by definition, after saccade offset. Note that the target never appeared at this screen location at any time during the experiment – any evidence of orientation tuning after the saccade must result from observers having attended to features at the pre-saccadic retinotopic location of the visual cue.

The spatio-temporal sensitivity kernels at the retinotopic trace location are shown in Fig S 2b. The modal saccade offset is shown as the dashed line at 50ms, at which time 62% of all saccades were complete, and 94% were complete by the end of the following time bin (+67ms). As shown in the upper plot of Fig S 2c, there is indeed tuning at the retinotopic location at saccade offset (“sac offset”), providing evidence for an attentional trace even though observers were never required to respond to a target at this location. This tuning curve, however, is relatively weak, and did not reach statistical significance. The attentional trace dissipated quickly, and within 34 ms of saccade offset the tuning curve is flat (“post-sac”, bottom plot of Fig S 2c).

**Fig S 2.**
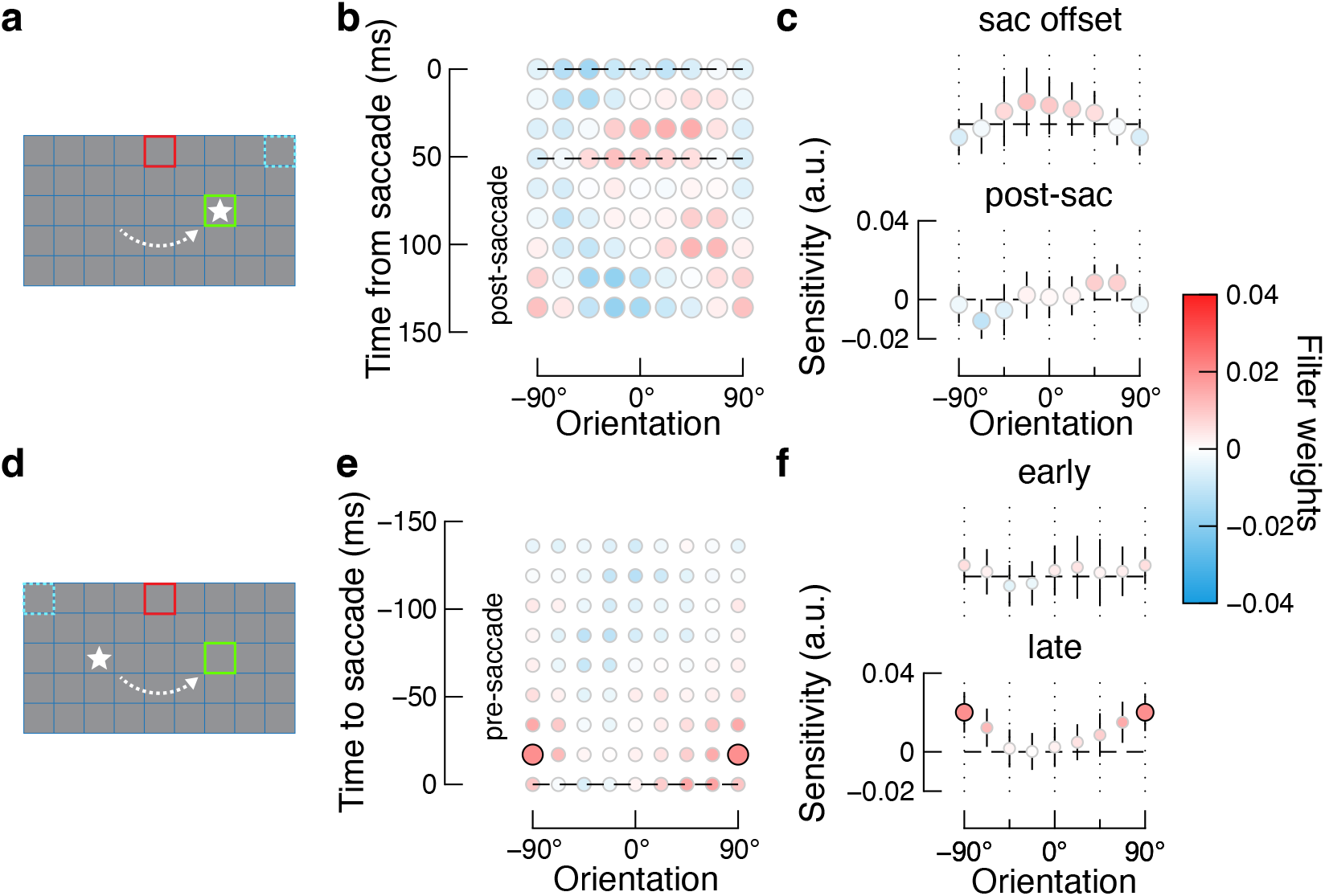
Testing for orientation selectivity at non-cued locations. a) The retinotopic trace location. As shown by the dotted blue square, the retinotopic trace location is the post-saccadic screen location that corresponds to the pre-saccadic retinal position of the cue. Other display features are described in Figure 1. b) Spatio-temporal sensitivity kernels at the retinotopic trace location. In spite of targets never appearing at this location, we find qualitative evidence for orientation selectivity. The colour scale for filter weights is the same as in Figure 5. c) Sensitivity kernels at the retinotopic trace location for two time slices after saccade onset. Although we find orientation selectivity when the saccade ends (sac offset, top), there is no selectivity 50ms later (post-sac, bottom), suggesting the retinotopic attentional trace is short-lived in time. d) The predictively remapped location. As shown by the dotted blue square, the remapped location is the pre-saccadic screen location that corresponds to the post-saccadic retinal position of the cue. e) Spatio-temporal sensitivity kernels at the remapped location. There is no clear evidence of orientation selectivity at any time point prior to the saccade. f) Sensitivity kernels at the remapped location for two time slices before saccade onset suggest no increase in perceptual benefits of attentional remapping in the period leading up to saccade onset.

### Predictive remapping of attention

To maintain attentional focus on the cued perceptual target location across saccades, a putative active remapping process must re-allocate attention from the cue’s pre-saccadic retinotopic location to its post-saccadic retinotopic location. The results of several studies suggest that such remapping begins before the eye movement ^9,22,45^. Within approximately 50ms *prior* to saccade onset, attentional performance benefits have been observed at a location that corresponds to a *post*-saccadic attentional cue location, the so-called predicted *remapped location* (Fig S 2d). We tested for such predictive remapping of attention by quantifying orientation tuning at the remapped location prior to saccade onset. As with the retinotopic trace location, the target never appeared at the remapped location. We are therefore uniquely able to test for predictive remapping of attention without presenting a perceptual probe at the remapped location, which might otherwise bias observers’ attention (as in previous studies ^42^).

The spatio-temporal sensitivity kernels at the remapped location are shown in Fig S 2e. There is no clear evidence for orientation tuning. However, predictive remapping of attention is thought to evolve in time such that attention benefits are greater immediately prior to the saccade than earlier in the pre-saccadic interval. We therefore show tuning curves at two different points in time in Fig S 2f. The “early” plot shows tuning 50ms prior to saccade onset, when any predictive remapping of attention should be weak or absent. The “late” plot shows tuning within 17ms prior to saccade onset, when the effects of predictive remapping should be strongest. There is a single significant tuning kernel in the late period, but the tuning is to features that are orthogonal to the target orientation. We suspect this is a spurious result; it cannot be predictive remapping because such tuning to non-target orientations would presumably be maladaptive.

## Methods

### Overview

The overarching aim of the study was to determine how saccadic eye movements and visual spatial attention are coordinated. We therefore had observers engage in a challenging visual discrimination task at an attended location off-fixation, and then investigated whether performance was influenced by the spatial and temporal dynamics of concurrently executed, goal-directed saccades (Figure 1). The visual discrimination task involved participants identifying a “dark” or “light” target bar embedded in dynamic visual noise. This design allowed investigation of attentional allocation at arbitrary screen locations, but without having to explicitly motivate observers to attend to multiple target locations, which might have biased the results of previous studies ^42^.

### Participants

Seven participants each completed approximately eleven separate testing sessions, each lasting up to 60 minutes in duration. All had normal or corrected-to-normal vision. Participants (five female) were aged between 20 and 36 years (M = 25 years). Five participants were naïve to the purpose of the experiment, and two were authors (WJH and IS). Ethics clearance was approved by the School of Psychology at The University of Queensland (ethics approval number: 18-PSYCH-4-43). Participation was voluntary, and all participants read an information sheet and signed a consent form.

### Psychophysical and eye tracking set-up

The experiment was programmed in MATLAB (The MathWorks, Inc., Natick, MA) with the Psychophysics Toolbox ^46,47^ and EyeLink Toolbox ^48^. All stimuli were presented on a 22-inch LED monitor (1080 x 1920 pixels; 120Hz refresh rate). An Eyelink 1000 was used to monitor participants’ eye position while they undertook the task. The position of participants’ right eye was sampled at a rate of 500Hz. The native eye-tracker 9-point calibration and validation procedure was performed at the start of each session and as required following breaks. Participants sat with their head stabilised on a chin rest positioned 57cm away from the monitor to limit head movements and to ensure accurate eye tracking. Perceptual reports were made via key press on a standard keyboard. The task was completed in a darkened and sound attenuated booth, and the experimenter monitored participants via an external monitor located outside the testing booth.

### Stimuli

A blue 13 x 13 grid was presented on a uniform grey background (luminance = 59 cd/m^2^). The width of each cell within the grid was 2.3° of visual angle. On each trial, two grid locations were coloured to indicate the required fixation location, and the cued target location. The fixation location was indicated by a green box, and the cued perceptual target location was indicated by a red box (for examples, see Figure 1). As described below, the green box shifted four grid locations either horizontally or vertically within a trial, whereas the red box was static within a trial but could shift across trials. The perceptual target location was always located midway between the start and end point of the required saccade, and was offset by two grid locations such that its centre had an eccentricity of 6.4° from the midpoint of the saccade start- and end-point in a trial. The luminance of the red and green boxes was perceptually equated.

Dynamic noise was presented within all grid locations during each trial. The noise consisted of a 9 x 9 array of spots, henceforth called “super-pixels” (the diameter of a super-pixel was approximately 10 pixels). The contrast of each super-pixel was independently drawn from a Gaussian distribution with a standard deviation of 12.5% contrast, updating at a rate of 60Hz. The perceptual target was a luminance increment or decrement added to one line of super-pixels at the cued target location. The centre of the target was the midpoint of the cued box, and its orientation was horizontal for vertical saccades, and vertical for horizontal saccades. We changed the target orientation according to saccade direction because pilot testing revealed that thresholds were elevated when the target orientation was parallel to saccade direction. The target thus appeared as either a “light” or “dark” modulated stripe orthogonal in orientation to the required saccade vector. Target duration was 17ms, and its amplitude was adjusted according to a staircase procedure, as described in detail below.

### Experimental procedure

Each participant was instructed that the aim of the experiment was to make saccades as quickly and accurately as possible while covertly attending to a target location and performing the visual discrimination task as accurately as possible. At the start of a trial, one grid location was outlined with a green box, indicating the required initial fixation position. Fixation compliance was verified online to ensure that participants were looking within 2° of the centre of the green box for 200 ms before the trial commenced. Concurrently with the fixation stimulus, one grid location was outlined in red to indicate the perceptual target location. The indicated target location was 100% predictive to motivate participants to covertly attend to this location continuously during the trial. After correct fixation had been verified, the green box disappeared and was immediately displayed at a new location within the grid to indicate the saccade goal. At the same time as the presentation of the saccade cue, dynamic noise was displayed across the display. Participants were instructed that the perceptual target could appear during any frame of the dynamic noise and that it would always appear within the red target square. After completion of the saccade and at the offset of the dynamic noise, participants reported whether the target was dark or light by pressing the left or right arrow keys, respectively. A break was enforced every 300 trials, and each session of the experimental task consisted of two blocks containing 600 trials each.

All possible fixation locations were constrained to fall within the central 9 x 9 portion of the grid, which ensured that the perceptual target location could always fall on either side of the saccade vector. The required saccade vectors were selected pseudo-randomly to ensure the saccade target never exceeded the limits of the inner 9 x 9 grid, but we ensured there were an equal number of vertical and horizontal eye movement trials within each block. The cued target location could appear two grid positions above or below horizontal saccade vectors, and likewise to the left or right of vertical saccade vectors, all with an equal probability. The target location was always positioned equidistant between the fixation point and the saccade landing location.

### Adaptive staircase and task practice

We controlled the amplitude of the target using an adaptive staircase to create sufficient perceptual difficulty to force observers to allocate attention to the cued location, as well as to ensure the dynamic noise influenced performance. We used a custom two-down, one-up staircase so that each observer’s accuracy was held at approximately 76% (sensitivity: d’ = 1 for two alternative forced choice task). Following two consecutive correct answers, the amplitude decreased by .90, and following one incorrect response, the amplitude increased by 1.1. Due to the relative difficulty of the task, each participant’s entire first session was considered as practice. In this session they completed three blocks of trials. First, they performed the perceptual task at the fovea, with no eye movements. Next, they performed the perceptual task in peripheral vision (i.e., while attending covertly to a location 6.4° from fixation) again without eye movements. Finally, participants practiced the full task with eye movements and perceptual reports.

## Analyses

### Eye data screening

Because our research question depended on observers correctly performing the task as described above, we adopted strict criteria to screen trials in which observers’ eye movements deviated from instructions. Trials were excluded from the analysis if saccadic latency was less than 100 ms or greater than 500 ms. Additionally, all trials in which participants broke fixation and looked toward the red perceptual target box were excluded. Trials were also excluded if the saccade landed more than 2.3° from the saccade goal (i.e., 25% of the required saccade amplitude), or if a blink was registered during a trial. In total, we excluded approximately 30,000 trials out of 84,000. As shown in Figure 2c, we nonetheless retained thousands of trials per participant, and hundreds of trials per target-saccade time bin.

### Behavioural performance

We quantified behavioural performance as observers’ proportion correct responses as well as contrast sensitivity derived from psychometric functions fit to responses across contrast levels. To investigate the time course of changes in perceptual performance across the saccade, the temporal dynamics of each trial were normalised with respect to saccade onset and offset: trials were binned according to the target onset relative to the onset of the saccade. The onset of the saccade was determined using the eye-tracker’s native saccade detection algorithm. Bins were 17ms in width.

Proportion correct was calculated as the average of correct responses within each time bin, separately for each observer (Figure 3a). To model contrast sensitivity, we fit a multilevel generalised linear model (GLMM), which allowed us to enter data from all participants and target-saccade asynchronies into a single model. The GLMM predicts perceptual reports from a linear combination of predictors:

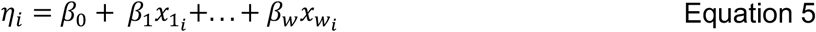

Where *η* is the sum of weighted linear predictors, *β*_*w*_ are the beta weights, and *x*_!_ are the predictors (i.e., the fixed effects) of the *i*th trial. The model was designed to capture any differences in sensitivity between negative and positive contrasts, as well as changes in sensitivity before and after eye movement onset. We therefore included three predictors (plus a bias term, *β*_0_): target contrast, the sign of the contrast (where negative values indicate the target amplitude was a luminance decrement relative to the mid-grey background), and whether the time interval was pre-or post-saccadic onset. We also included all interactions between predictors. The sum of these weighted predictors was then passed through a probit link function to estimate proportion of “light” responses:

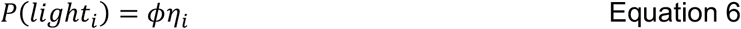

Where *ϕ* is the normal integral function. We implemented the model as a GLMM to partially pool coefficient estimates across observers and time points ^49,50^. We modelled each observer’s predictor weights and each time-point’s predictor weights as having come from independent population distributions with mean *μ* and variance *σ*^2^:

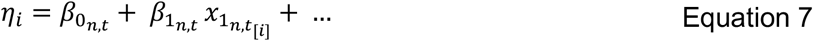

Where

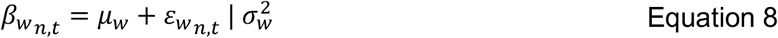

Here, 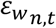 is the offset of the predictor for each observer, *n*, and each time-point, *t*, relative to the parameter’s mean, *μ*_*w*_, contingent on the estimated population variance, 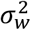. The effect of partial pooling in the GLMM is to pull extreme values toward the mean, to an extent that is proportional to the estimated variance in each measurement. Such pooling provides a more conservative estimate of the influence of predictors than in a standard GLM, but these differences are likely to be relatively minor in our data because of the large number of trials, and therefore precise estimates, for each observer and time point. The model was fit using MATLAB’s fitglme() function.

### Dynamic orientation selectivity across saccades

We used reverse correlation to quantify the influence of visual noise on observers’ perceptual reports, allowing us to recover the spatio-temporal characteristics of their attentional allocation across saccades. Whereas reverse correlation is commonly performed on each noise pixel to construct a classification image ^18,e.g. 23^, we instead applied it to the contrast energy of various filters defined by their orientation and spatial frequency ^51^. Note that this analysis was performed on the target-free noise; we were only interested in the effects of visual noise and not the target itself (which would otherwise contaminate the correlational analysis). We included in the analysis only those trials that were not excluded in the screening procedures outlined above. We describe the analysis as performed at the cued perceptual target location; other analyses were identical but performed on noise at different screen positions as described in text.

We first aligned the visual noise to a common temporal sequence across trials by time-locking the noise according to saccade onset. The noise frame during which saccade onset was detected was assigned the temporal position of 0ms, equivalent to the 0ms target-onset asynchrony condition in the behavioural data. To ensure that differences in saccade durations could not contaminate our estimates of peri-saccadic tuning, we further assigned the noise during which the saccade ended as time +50ms. The 67ms time bin therefore included the first noise frame post-saccadic offset for all trials. Similarly, the -17ms time bin included the final frame before saccade onset for all trials. We also transformed the relative spatial organisation of the noise such that all trials were spatially aligned relative to a rightward saccade and with the perceptual target location above the saccade vector, as shown in Figure 1b and c.

We then measured the response of a bank of two-dimensional Gabor filters that varied in orientation and spatial frequency to each noise frame. This is equivalent to measuring the output of a population of neurons whose receptive fields were centred on the target location throughout the duration of the trial at a temporal resolution of 60Hz. Gabors varied in orientation from 0° to 180° in 22.5° steps, ranged in spatial frequency from 1 cycle per image to 4.5 cycles per image (the Nyquist limit given the noise dimensions), and had a Gaussian envelope with a standard deviation of 1.5 super-pixels. All filters were phase-matched to the target (phase = 90°; even-symmetric filters). The response of each filter at each time point is given by:

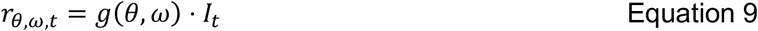

Where *r*_*θ ωt*_ is the output of a filter, *g*, with orientation *θ* and spatial frequency *ω*, in response to the noise, *I*, at a time *t* on a given trial, and the ⋅ operator indicates the dot (or inner) product. We then applied divisive normalisation ^52^ to the filter responses within each spatial frequency band, for each noise frame:

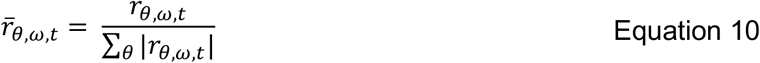

To ensure all sensitivity kernels were comparable, we further normalised filter responses to have a mean of zero and standard deviation of 1. Finally, we used a series of GLMMs to predict the probability an observer would report the target as being relatively “light” from 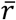from each filter combination and at each time point:

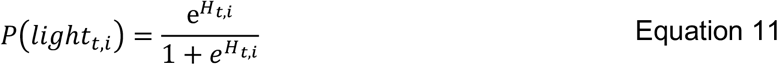

And

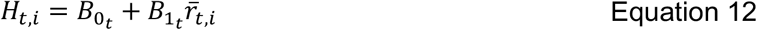

Where *H* is the linear predictor at time point *t* on trial *i*, 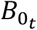 is a bias term, and 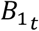 is the correlation between a filter’s response and the perceptual report at time *t*. As with the behavioural data, we fit this model using MATLABs fitglme() function and modelled participants as a Gaussian-distributed random effect. Prior to the calculation of the filter responses, we applied a small amount of spatio-temporal smoothing to mimic the integration window of the visual system ^23^. Smoothing was performed using a three-dimensional Gaussian kernel, with standard deviations of 0.1, 0.1, and 20, corresponding to x, y, and time coordinates, respectively. Modifications to the smoothing kernel dimensions, the number of oriented filters, or the peak and width of spatial frequency bands did not alter the results.

The modelling procedure described above results in a three-dimensional kernel, *K*_*θ ωt*_, that describes the average observer’s attentional selectivity in orientation, spatial frequency, and time. The full set of kernels within *K* _*θ ωt*_ is shown in Supp Fig S 1.

### An ideal observer model that allocates attention in real world coordinates

The critical aspects of the ideal observer are described in the main text and in Equations 1 – 4. These equations show how the real-world position of the cued target location can be estimated from the gaze position and the retinal position of the stimulus, and how the ideal observer makes a perceptual decision by allocating attention in the form of a template at the estimated target location.

For each observer, the model completed the same number of trials as included in the analyses described in previous sections. On each trial, the model executed a saccade of variable duration, as per the real observers. We parameterised each observer’s saccadic latencies as a gamma distribution, and generated saccadic durations by drawing gamma-distributed random variables with the shape and scale parameters for each participant. The spatial dynamics of a saccade were modelled as a half cosine function, smoothly transitioning between two points separated by the equivalent of 9° throughout the duration of the saccade:

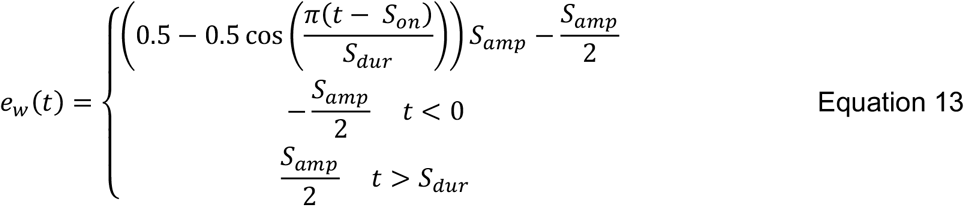

Where *S*_0*n*_ is relative saccade onset time, *S*_*dur*_, is saccadic duration, and *S*_*amp*_ is saccadic amplitude. We set *S*_0 *n*_ to 0ms, *S*_*amp*_ to 9°, and *S*_*dur*_ as per the gamma distributions described above. An example of a saccade of 50ms duration is shown in Figure 6a.

On each simulated frame of a trial, the model observer applied the attentional template at the estimated real-world location of the cue, as described in Equation 1, and calculated and reported the sign of the response of the template as per Equations 2 – 4. The visual noise presented at the cued target location (not the estimated location) was then analysed as per the reverse correlational analyses described above. Note that no free parameters were used to determine how the observer model made eye movements.

To estimate the cued targets’ real-world location, the observer model combined information about the retinal position of the object and gaze position as estimated by gain-field neurons. To simplify these calculations, our observer was designed to execute a saccade from 4.5° to the left of the midline of the screen to 4.5° to the right of the midline of the screen, positioning the cued target location directly on the vertical midline. Under these circumstances and as shown in Figure 6a, the retinal position of the cued target location is simply half of the negative of the current gaze position. Rather than modelling the spikes of idealised gain-field neurons and then applying the same Bayesian decoder as Morris and Krekelberg ^13^, we instead modelled only the decoded gaze position signal. Morris and Krekelberg found that the decoded gaze position lags the true eye position by approximately 45ms. We therefore modelled the estimated gaze position as being equivalent to the saccade trajectory, but with a time course that was delayed in time by some constant, which we refer to as *gain field delay*. This was the single free parameter in our observer model. To estimate its value, we simulated a full set of data for each observer with gain field delays ranging from 0ms – 75ms, in 5ms steps, performed the reverse correlation on each set of data, and then fit the kernel estimates of the observer model to the empirical data using a GLM per observer per gain field delay. We then found the gain field delay that minimised the deviance of the model fit for each observer separately. Because of the stochastic processes involved in the simulations, such as the variable saccadic durations and dynamic visual noise, we repeated this process 50 times, finding the global minima for each observer and the group. Due to the volume of data generated during this procedure, as well as the high number of reverse correlational analyses performed, the full modelling procedure took approximately 48 hours to complete on a 2018 iMac.

## References

1. Irwin, D.E. (1996). Integrating information across saccadic eye movements. Curr. Dir. Psychol. Sci. 5, 94–100.

2. Dorr, M., Martinetz, T., Gegenfurtner, K.R., and Barth, E. (2010). Variability of eye movements when viewing dynamic natural scenes. J. Vis. 10, 28. 10.1167/10.10.28.

3. Remington, R.W. (1980). Attention and saccadic eye movements. J. Exp. Psychol. Hum. Percept. Perform. 6, 726–744.

4. Dorr, M., and Bex, P.J. (2013). Peri-Saccadic Natural Vision. J. Neurosci. 33, 1211–1217. 10.1523/JNEUROSCI.4344-12.2013.

5. Burr, D.C., Morrone, M.C., and Ross, J. (1994). Selective suppression of the magnocellular visual pathway during saccadic eye movements. Nature 371, 511–513. 10.1038/371511a0.

6. Diamond, M.R., Ross, J., and Morrone, M.C. (2000). Extraretinal control of saccadic suppression. J. Neurosci. 20, 3449–3455.

7. Volkmann, F.C., Riggs, L.A., White, K.D., and Moore, R.K. (1978). Contrast sensitivity during saccadic eye movements. Vision Res. 18, 1193–1199. 10.1016/0042-6989(78)90104-9.

8. Cavanagh, P., Hunt, A.R., Afraz, A., and Rolfs, M. (2010). Visual stability based on remapping of attention pointers. Trends Cogn. Sci. 14, 147–153. 10.1016/j.tics.2010.01.007.

9. Rolfs, M., Jonikaitis, D., Deubel, H., and Cavanagh, P. (2011). Predictive remapping of attention across eye movements. Nat. Neurosci. 14, 252–256. 10.1038/nn.2711.

10. Colby, C.L., and Goldberg, M.E. (1999). Space and attention in parietal cortex. Annu. Rev. Neurosci. 22, 319–349.

11. Melcher, D., and Colby, C.L. (2008). Trans-saccadic perception. Trends Cogn. Sci. 12, 466–473. 10.1016/j.tics.2008.09.003.

12. Bays, P., and Husain, M. (2007). Spatial remapping of the visual world across saccades. Neuroreport 18, 1207.

13. Morris, A.P., and Krekelberg, B. (2019). A Stable Visual World in Primate Primary Visual Cortex. Curr. Biol. 29, 1471-1480.e6. 10.1016/j.cub.2019.03.069.

14. Hubel, D.H., and Wiesel, T.N. (1959). Receptive fields of single neurones in the cat’s striate cortex. J. Physiol. 148, 574–591.

15. Campbell, F.W., and Robson, J.G. (1968). Application of Fourier analysis to the visibility of gratings. J. Physiol. 197, 551–566.

16. Weyand, T.G., and Malpeli, J.G. (1993). Responses of neurons in primary visual cortex are modulated by eye position. J. Neurophysiol. 69, 2258–2260. 10.1152/jn.1993.69.6.2258.

17. Trotter, Y., and Celebrini, S. (1999). Gaze direction controls response gain in primary visual-cortex neurons. Nature 398, 239–242. 10.1038/18444.

18. Harrison, W.J., and Rideaux, R. (2019). Voluntary control of illusory contour formation. Atten. Percept. Psychophys. 81, 1522–1531. 10.3758/s13414-019-01678-8.

19. Mareschal, I., Dakin, S.C., and Bex, P.J. (2006). Dynamic properties of orientation discrimination assessed by using classification images. Proc. Natl. Acad. Sci. U. S. A. 103, 5131–5136. 10.1073/pnas.0507259103.

20. Ringach, D.L., Hawken, M.J., and Shapley, R. (1997). Dynamics of orientation tuning in macaque primary visual cortex. Nature 387, 281–284. 10.1038/387281a0.

21. Golomb, J.D., Chun, M.M., and Mazer, J.A. (2008). The native coordinate system of spatial attention is retinotopic. J. Neurosci. 28, 10654–10662. 10.1523/JNEUROSCI.2525-08.2008.

22. Jonikaitis, D., Szinte, M., Rolfs, M., and Cavanagh, P. (2013). Allocation of attention across saccades. J. Neurophysiol. 109, 1425–1434. 10.1152/jn.00656.2012.

23. Neri, P., and Heeger, D.J. (2002). Spatiotemporal mechanisms for detecting and identifying image features in human vision. Nat. Neurosci. 5, 812–816. 10.1038/nn886.

24. Panichi, M., Burr, D., Morrone, M.C., and Baldassi, S. (2012). Spatiotemporal dynamics of perisaccadic remapping in humans revealed by classification images. J. Vis. 12, 1–15. 10.1167/12.4.11.

25. Golomb, J.D. (2019). Remapping locations and features across saccades: a dual-spotlight theory of attentional updating. Curr. Opin. Psychol. 29, 211–218. 10.1016/j.copsyc.2019.03.018.

26. Geisler, W.S. (2018). Psychometric functions of uncertain template matching observers. J. Vis. 18, 1–10. 10.1167/18.2.1.

27. Müller, H.J., and Rabbitt, P.M. (19891001). Reflexive and voluntary orienting of visual attention: Time course of activation and resistance to interruption. J. Exp. Psychol. Hum. Percept. Perform. 15, 315. 10.1037/0096-1523.15.2.315.

28. Rolfs, M., and Carrasco, M. (2012). Rapid Simultaneous Enhancement of Visual Sensitivity and Perceived Contrast during Saccade Preparation. J. Neurosci. 32, 13744–13752. 10.1523/JNEUROSCI.2676-12.2012.

29. Yao, T., Ketkar, M., Treue, S., and Krishna, B.S. (2016). Visual attention is available at a task-relevant location rapidly after a saccade. eLife 5, e18009. 10.7554/eLife.18009.

30. Harrison, W.J., and Bex, P.J. (2014). Integrating retinotopic features in spatiotopic coordinates. J. Neurosci. 34, 7351–7360. 10.1523/JNEUROSCI.5252-13.2014.

31. Collins, T., Rolfs, M., Deubel, H., and Cavanagh, P. (2009). Post-saccadic location judgments reveal remapping of saccade targets to non-foveal locations. J. Vis. 9, 29.1-9. 10.1167/9.5.29.

32. Marino, A.C., and Mazer, J.A. (2018). Saccades Trigger Predictive Updating of Attentional Topography in Area V4. Neuron 98, 429-438.e4. 10.1016/j.neuron.2018.03.020.

33. Bergelt, J., and Hamker, F.H. (2019). Spatial updating of attention across eye movements: A neuro-computational approach. J. Vis. 19, 10–10. 10.1167/19.7.10.

34. Duhamel, J.R., Colby, C.L., and Goldberg, M.E. (1992). The updating of the representation of visual space in parietal cortex by intended eye movements. Science 255, 90–92.

35. Võ, M.L.-H., Aizenman, A.M., and Wolfe, J.M. (2016). You think you know where you looked? You better look again. J. Exp. Psychol. Hum. Percept. Perform. 42, 1477–1481. 10.1037/xhp0000264.

36. Clarke, A.D.F., Mahon, A., Irvine, A., and Hunt, A.R. (2016). People are unable to recognize or report on their own eye movements. Q. J. Exp. Psychol., 1–56. 10.1080/17470218.2016.1231208.

37. Wurtz, R.H. (2008). Neuronal mechanisms of visual stability. Vision Res. 48, 2070–2089. 10.1016/j.visres.2008.03.021.

38. Bays, P.M., and Husain, M. (2008). Dynamic shifts of limited working memory resources in human vision. Science 321, 851–854. 10.1126/science.1158023.

39. Morris, A.P., Chambers, C.D., and Mattingley, J.B. (2007). Parietal stimulation destabilizes spatial updating across saccadic eye movements. Proc. Natl. Acad. Sci. U. S. A. 104, 9069–9074. 10.1073/pnas.0610508104.

40. Bottini, R., and Doeller, C.F. (2020). Knowledge Across Reference Frames: Cognitive Maps and Image Spaces. Trends Cogn. Sci. 24, 606–619. 10.1016/j.tics.2020.05.008.

41. Golomb, J.D., Pulido, V.Z., Albrecht, A.R., Chun, M.M., and Mazer, J.A. (2010). Robustness of the retinotopic attentional trace after eye movements. J. Vis. 10, 19.1-12. 10.1167/10.3.19.

42. Arkesteijn, K., Belopolsky, A.V., Smeets, J.B.J., and Donk, M. (2019). The Limits of Predictive Remapping of Attention Across Eye Movements. Front. Psychol. 10. 10.3389/fpsyg.2019.01146.

43. Deubel, H., and Schneider, W.X. (1996). Saccade target selection and object recognition: evidence for a common attentional mechanism. Vision Res. 36, 1827–1837.

44. Harrison, W.J., Mattingley, J.B., and Remington, R.W. (2013). Eye movement targets are released from visual crowding. J. Neurosci. 33, 2927–2933. 10.1523/JNEUROSCI.4172-12.2013.

45. Szinte, M., Jonikaitis, D., Rangelov, D., and Deubel, H. (2018). Pre-saccadic remapping relies on dynamics of spatial attention. eLife 7, e37598. 10.7554/eLife.37598.

46. Brainard, D.H. (1997). The Psychophysics Toolbox. Spat. Vis. 10, 433–436.

47. Pelli, D.G. (1997). The VideoToolbox software for visual psychophysics: Transforming numbers into movies. Spat. Vis. 10, 437–442.

48. Cornelissen, F.W., Peters, E.M., and Palmer, J. (2002). The Eyelink Toolbox: eye tracking with MATLAB and the Psychophysics Toolbox. Behav. Res. Methods Instrum. Comput. 34, 613–617.

49. Gelman, A., and Hill, J. (2007). Data Analysis Using Regression and Multilevel/Hierarchical Models (Cambridge University Press).

50. Rideaux, R., West, R.K., Wallis, T.S.A., Bex, P.J., Mattingley, J.B., and Harrison, W.J. (2022). Spatial structure, phase, and the contrast of natural images. J. Vis. 22, 1–19. 10.1167/jov.22.1.4.

51. Li, H.-H., Barbot, A., and Carrasco, M. (2016). Saccade Preparation Reshapes Sensory Tuning. Curr. Biol. 26, 1564–1570. 10.1016/j.cub.2016.04.028.

52. Carandini, M., and Heeger, D.J. (2012). Normalization as a canonical neural computation. Nat. Rev. Neurosci. 13, 51–62. 10.1038/nrn3136.

